# Tumor protein D52 (TPD52) affects cancer cell metabolism by negatively regulating AMPK

**DOI:** 10.1101/2021.03.09.434668

**Authors:** Yali Chen, Changmin Peng, Wei Tan, Jia Yu, Jacqueline Zayas, Yihan Peng, Zhenkun Lou, Huadong Pei, Liewei Wang

## Abstract

The AMP-activated protein kinase (AMPK) is a central regulator of energy homeostasis, with deregulation leading to cancer and other diseases. When intracellular ATP levels decrease during energy stress, AMPK is phosphorylated and activated through AMP binding. However, how this pathway is dysregulated in cancer remains unclear. Here, we find that tumor protein D52 (TPD52), initially identified to be overexpressed in many human cancers, forms a stable complex with AMPK in cancer cells. TPD52 directly interacts with AMPKα and inhibits AMPKα kinase activity *in vitro* and *in vivo*. We generated TPD52 transgenic mice, and found that overexpression of TPD52 leads to AMPK inhibition and multiple metabolic defects in mice. Together, our results shed new light on AMPK regulation and on our understanding of the etiology of cancers with TPD52 overexpression.

## Introduction

AMPK is one of the most important molecular energy sensors in eukaryotic cells (1,2). AMPK is a 5’-AMP-activated protein kinase, which is an evolutionarily conserved heterotrimer formed by a catalytic α subunit and two regulatory subunits (β and γ) (3–6). The γ subunit regulates AMPK ability to sense the AMP/ATP ratio. The γ subunit contains four cystathionine beta-synthase (CBS) domains, which are binding sites for AMP/ADP/ATP (7). When cells have an insufficient energy supply, the γ subunit of AMPK binds to AMP, thereby sensing the increased AMP/ATP ratio. Consequently, the AMP-bound γ subunit leads to a major conformational change in the AMPK heterotrimer complex, enabling the exposure of the catalytic pocket of α subunit and activation of AMPK kinase (8–10). AMPK activation requires the phosphorylation of Thr172 in α subunit by the upstream kinase liver kinase B1 (LKB1) (11–14). AMPKα can also be phosphorylated at Thr172 in response to calcium flux by CaMKKβ (15–17).

AMPK is able to control a wide range of metabolic processes that connect cellular metabolism with energy availability (18–20). AMPK controls glucose uptake and fatty acid oxidation in muscle, fatty acid synthesis and gluconeogenesis in liver, and the regulation of food intake in hypothalamus (21–23). AMPK transmits glucose shortage or energy crisis signal to downstream targets via phosphorylation events (24,25). For example, AMPK phosphorylates acetyl-CoA carboxylase 1 (ACC1) and sterol regulatory element-binding protein 1c (SREBP1c) to suppress lipid and cholesterol synthesis (26,27); AMPK phosphorylates ULK1 to modulate autophagy, which helps cells to adapt to cellular energetic status (28,29); AMPK phosphorylates TSC2 to inhibit Rheb GTP loading, and renders mTORC1 inactivation (30).

The LKB1-AMPK pathway is a central regulator of energy metabolism, and dysregulation of this pathway has been implicated in cancers, including breast cancer (31–33). Indirect AMPK activators such as metformin have shown beneficial effect in breast cancer prevention and treatment (34,35). However, other than LKB1 mutations, how this pathway might be dysregulated in cancer remains unclear. Therefore, it will be important to identify additional novel regulators in the LKB1-AMPK signaling pathway that may be involved in developing both acquired and inherited forms of cell metabolism abnormalities.

In the present study, we identified TPD52 as a novel regulator of the LKB1-AMPK pathway. TPD52 is known to be involved in the regulation of vesicle trafficking and exocytotic secretion (36–38). In addition, the 8q21 locus containing the *Tpd52* gene is frequently amplified in tumors, including breast (39–42). However, the exact mechanisms of TPD52 in cancer metabolism remain unclear. Our results demonstrated that TPD52 affects cancer cell metabolism by negatively regulating AMPK.

## Materials and methods

### Cell culture

HEK-293T cells were maintained in Dulbecco’s modified Eagle’s medium (DMEM) supplied with 10% fetal bovine serum (FBS) and penicillin/streptomycin (P/S) at 37°C with 5% CO_2_. Human breast carcinoma cell lines (SK-BR-3, MCF-7, MDA-MB-231, T-47D) were cultured in GIBCO® RPMI 1640 (Life technologies) medium supplemented with 10% FBS (Life technologies), 6 mM L-glutamine (Life technologies), and 10 μg/ml insulin (Sigma-Aldrich) (for MCF-7) in a humidified atmosphere containing 5% CO_2_ at 37°C.

### Tpd52 transgenic mice

*Tpd52* transgenic founder mice were generated by Cyagen Biosciences Inc (Suzhou, China). Briefly, the PiggyBac transposon gene expression vector for mouse *Tpd52* gene (*mTpd52*) was created by cloning of the CAG promoter plus the open reading frame (ORF) of *mTpd52* gene with an HA-tag into the Cyagen basic PiggyBac (PB) vector and then verified by sequencing. The linearized vector DNA of pPB-CAG-*mTpd52* ORF-HA was purified using a QIAquick Gel extraction kit (Qiagen, Chatsworth, CA, USA) and adjusted to a final concentration of 3 ng/μl in Tris-EDTA buffer for pronucleus injection. The fertilized one-cell eggs from the superovulated female C57BL/6 mice were injected and implanted into the oviducts of pseudopregnant female C57BL/6 mice. The positive founders were screened by genotyping using primers (forward 5’-CTGGTTATTGTGCTGTCTCATCAT-3’ and reverse 5’-TCAGCAGACCAACGTTCTGTG-3’) with a product of 214-bp fragment. The identified positive founders were then bred with wild-type C57BL/6 mice for generation of the *mTpd52* transgenic mice.

### Constructs, antibody and siRNA

TPD52 full length cDNA or TPD52 deletion mutants were cloned into a modified pIRES2-N-FLAG-S-tag vector to generate mammalian expression constructs with N terminal FLAG and S-protein tag. HA-tagged TPD52 expression constructs were generated by cloning TPD52 cDNA into pCMV-HA vector. TPD52 cDNA was also cloned into pGEX-4T-2 to generate GST fusion protein expression plasmids. His-tagged AMPKα, β and γ prokaryotic expression constructs were generated from pET-28a. All the mutations used in this study were generated using site-directed mutagenesis kit, and were verified by sequencing.

The following antibodies were used in our study: phospho-AMPKα (2535S, WB: 1:1000 dilution; Cell Signaling Technology), AMPKα (5832S, WB: 1:1000 dilution; Cell Signaling Technology), AMPKβ (4150S, WB: 1:1000 dilution; Cell Signaling Technology), AMPKγ (4187S, WB: 1:1000 dilution; Cell Signaling Technology), phospho-ACC1 (3661S, WB: 1:1000 dilution; Cell Signaling Technology), ACC1 (3662S, WB: 1:1000 dilution; Cell Signaling Technology), TPD52 (GTX115042, WB:1:400 dilution; GeneTex), Anti-His tag (sc8036, WB: 1:1000; Santa Cruz), Rabbit anti-FLAG (F7425, WB: 1:2500; Sigma-Aldrich), Mouse anti-FLAG (F3165, WB: 1:2500; Sigma-Aldrich), Mouse anti-HA (901501, WB: 1:2500; IF: 1:100; Biolegend).

Two sets of TPD52 siRNA duplexes were synthesized by Thermo Fisher Scientific. The targeted sequences were: 5’-GCGGAAACUUGGAAUCAAU-3’ (siTPD52-1), and 5’-GGAGAAGUCUUGAAUUCGG-3’ (siTPD52-2). Non-targeting siRNA (All-star Negative Control siRNA) was purchased from QIAGEN.

### Immunoprecipitation Assay

Cells transfected with indicated constructs were harvested and lysed in NETN buffer (10mM Tris-HCl [pH 8.0], 100mM NaCl, 1mM EDTA, and 0.5% NP-40) plus protease inhibitor (Roche) on ice for 30 min. Then cell lysates were subject to protein immunoprecipitations using indicated beads (For N-Flag-S-protein-tagged, FLAG M2 (Sigma-Aldrich) or S-protein beads (Millipore) were used. For HA-tagged, HA beads (Sigma-Aldrich) were used), After 8 hr rotation at 4°C, beads were washed with NETN buffer three times, and samples were eluted with 2× SDS loading buffer and were subject to immunoblot with indicated antibodies.

For endogenous IP, cell lysates were incubated with indicated antibodies at 4°C for 6 hr, and then were mixed with Protein A/G beads (Thermo Fisher Scientific) at 4°C for 4 hr. Beads were then washed with NETN buffer 3 times, and samples were eluted with 2× SDS loading buffer, and were analyzed by immunoblot with indicated antibodies.

### Protein purification

For GST-tagged protein purification, GST fusion proteins were expressed and purified from *E.coli* BL21 bacteria, and were then immobilized on Glutathione Sepharose 4B (GE healthcare) at 4°C overnight. Proteins were then eluted with 5 folds volume of Glutathione solution (Sigma-Aldrich), and were condensed using 10kD protein concentrators (Thermo Fisher Scientific). Proteins were resolved, and stocked in PBS containing 5% Glycerol at −80°C.

For His-tagged Protein purification. His-tagged proteins were expressed and purified from *E.coli* BL21 bacteria, and were immobilized on Ni-NTA (Thermo Fisher Scientific) at 4°C. Proteins were then eluted with 10 folds volume of 300mM imidazole, and were dialyzed and condensed using 10kD protein concentrators. Protein were resolved, and stocked in PBS containing 5% Glycerol at −80°C.

### GST pull down Assay

GST fusion proteins were purified from *E.coli* BL21 bacteria, and were immobilized on Glutathione Sepharose 4B at 4°C overnight. HEK293T cells transfected with indicated constructs were treated as indicated. Then cells were lysed in NETN buffer plus protease inhibitors and were incubated with the Sepharose beads immobilized with indicated proteins at 4°C for 8 hr. Sepharose beads were then washed with NETN buffer five times and were eluted with 2× SDS loading buffer. Samples were subject to immunoblot with indicated antibodies.

### Oil Red O staining

The intracellular lipid droplet contents of cultured cells were evaluated by Oil Red O staining. Lipids in mouse liver were extracted as described (43). Briefly, cells were washed with ice-cold PBS, fixed with 10% formalin for 60 min, and stained with Oil-Red-O working solution (1.8 mg/ml of Oil-Red-O in 6:4 isopropanol: water solution) for 60 min at 25°C. After staining, cells were washed with water to remove any remaining dye. For quantification of Oil-Red-O staining, the cell-retained dye was extracted by isopropanol and the content was measured spectrophotometrically at 500 nm.

### mRNA extract and qRT-PCR

Total mRNA was isolated from cells using PARIS™ Kit (Thermo Fisher Scientific). One microgram of total RNA was reverse transcribed with random hexamers using Superscript II reverse transcriptase (Invitrogen) according to the manufacturer’s protocol. Real-time PCR was performed on a Bio-Rad iCycler using IQ SYBR green (Bio-Rad) with the following primers: mouse *actin* primers (forward 5’-CGGTTCCGATGCCCTGAGGCTCTT-3’ and reverse 5’-CGTCACACTTCATGATGGAATTGA-3’); mouse *ldha* primers (forward 5’-TGTCTCCAGCAAAGACTACTGT-3’ and reverse 5’-GACTGTACTTGACAATGTTGGGA-3’); mouse *pdk1* primers (forward 5’-ACAAGGAGAGCTTCGGGGTGGATC-3’ and reverse 5’-CCACGTCGCAGTTTGGATTTATGC-3’; mouse *fas* primers (forward 5’-GCTGCGGAAACTTCAGGAAAT-3’ and reverse 5’-AGAGACGTGTCACTCCTGGACTT-3’); mouse *scd1* primers (forward 5’-CTGACCTGAAAGCCGAGAAG-3’ and reverse 5’-GCGTTGAGCACCAGAGTGTA-3’).

### Animal diet

WT and transgenic male mice at the age of 6 weeks were fed on a nonfat diet (NFD) or high fat diet (HFD, 60 kcal% Fat, D12492, QiFa Biotech., Shanghai) for 16 week. GTT assay was performed in 16-week old mice. Blood glucose level was performed in 20-week old mice. IHC and oil red O staining and qRT-PCR was performed with liver tissue samples from 22-week old mice. These procedures were approved by the Institutional Animal Care and Use Committee (IACUC).

### Liver histological and immunohistochemical analysis

Livers were fixed in 10% phosphate-buffered formalin acetate at 4°C overnight and embedded in paraffin wax. Paraffin sections (5μm) were cut and mounted on glass slides for hematoxylin and eosin (H&E) staining. Cryosections of livers were stained by Oil Red O and counterstained with hematoxylin to visualize the lipid droplets. Immunohistochemistry of liver sections was also performed to visualize phosphorylated ACC1, phosphorylated AMPK and TPD52.

### Blood glucose detection

Fast mice for 16 hr. Blood glucose levels were measured in tail vein blood samples using a glucometer. Values are expressed as mean ± SD (n = 6).

### IPGTT (glucose tolerance test)

For IPGTT, after 5 hr fasting, male mice were injected with 1g/kg bodyweight glucose intraperitoneally. Blood glucose was measured before the glucose injection and at 15, 30, 60 and 90 min post-injection. Values are expressed as mean ± SD (n = 6).

### Statistical Analysis

mRNA levels were determined by examining the RNA-seq values of tumors in the cancer genome atlas (TCGA). Based on the FPKM values of each gene, we excluded genes with low expression, i.e., less than FPKM 1. Survival data were determined by clinical data performed on tumors with matching RNA-seq data (https://portal.gdc.cancer.gov/repository, TCGA-BRCA, V13.0, 2018). A median FPKM cut-off (27.70765485) was chosen for dividing the patients into low- and high-TPD52 expression groups. Survival distributions were estimated by the Kaplan-Meier method, and the significance of differences between overall survival rates was ascertained using the log-rank test. Gene expression correlation between TPD52 and other metabolic genes were analyzed for statistical significance using Pearson correlation (R).

All the other statistical analysis was performed with data from three biological triplicates. Statistical analysis was performed by the Student’s *t*-test for two groups and by ANOVA for multiple groups. *P*<0.05 was considered significant.

## Results

### TPD52 interacts with AMPKα

TPD52 is known to be commonly overexpressed in breast cancer, but the physiological functions are not clear. To gain greater insight into the *in vivo* function of TPD52, we overexpressed TPD52 and purified the TPD52 complex from 293T cells using the sequential double-affinity tag purification technique described previously (44,45). The resulting TPD52 complex was subjected to tandem mass spectrometry. A number of known TPD52-associated proteins were co-purified, including TPD52L1, TPD52L2, SRPK1 and SRPK2 (Figure 1(a)). Interestingly, we also identified a series of peptides corresponding to the two subunits of the kinase, AMPK (AMPKα1, AMPKα2, AMPKβ1 and AMPKβ2). To further confirm this interaction, we overexpressed and purified His-tagged AMPKα1, AMPKα2, AMPKβ1, AMPKβ2 and AMPKγ1 from *E.coli*, and then tested their binding to GST-TPD52 *in vitro*. As shown in Figure 1(b), TPD52 directly bound to AMPK α1 and α2, but not the other AMPK subunits. We also performed a co-immunoprecipitation assay with anti-AMPKα antibody and found that endogenous AMPKα was associated with TPD52 in cells (Figure 1(c)). We also determined the interactions between TPD52 and AMPKα1 kinase domain (1-300 amino acids, AMPKαE) using a GST-pull-down assay (Figure 1(d)). We next investigated which regions of TPD52 might be responsible for the interaction with AMPKα by expressing full length TPD52 or a series of truncated mutants in HEK-293T cells. TPD52 deletion mutant D4 (deletion amino acid residues 1–61) abolished the binding of TPD52 with AMPKα (Figure 1(e)).

**Figure 1.**
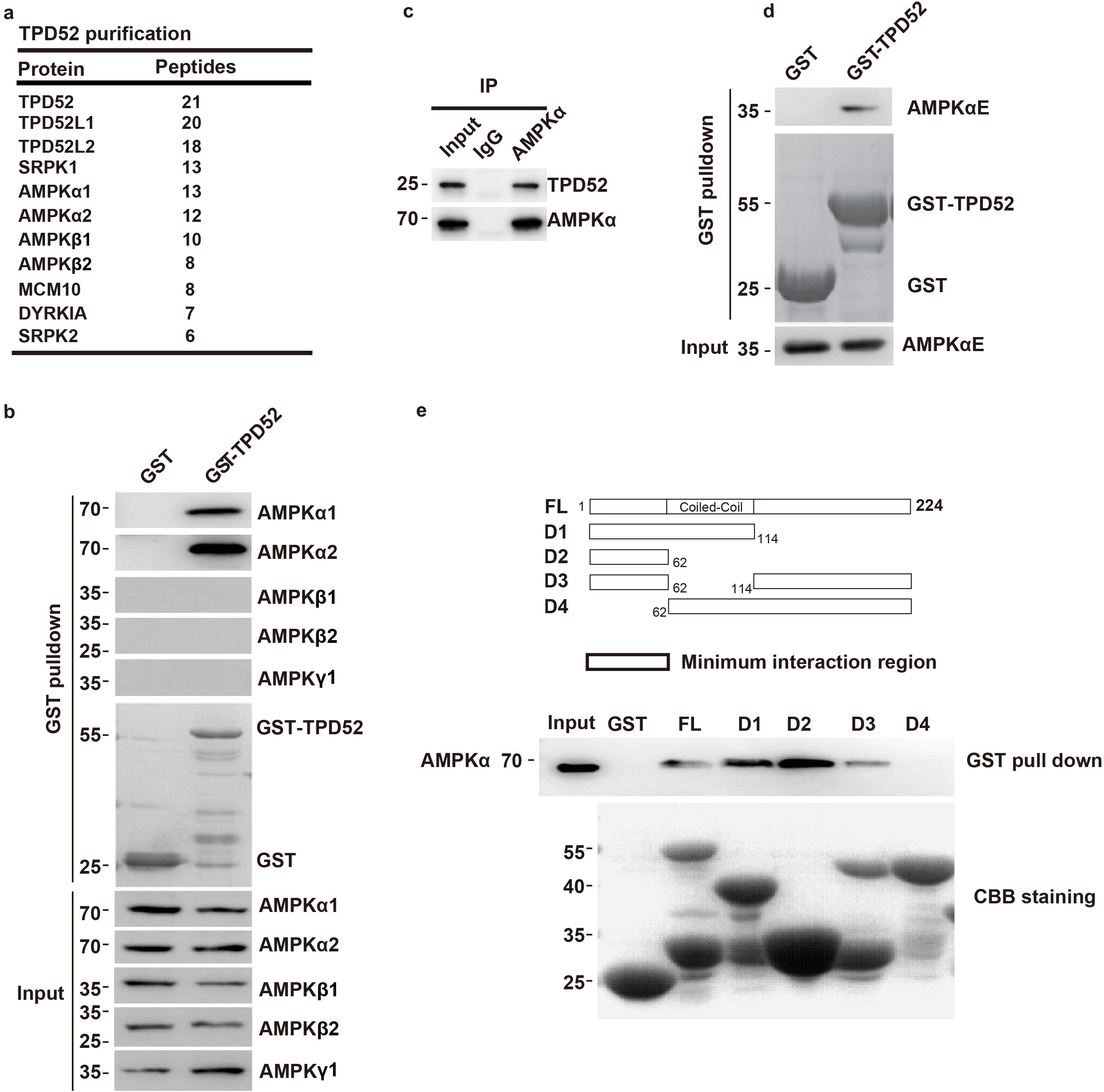
TPD52 interacts with AMPKα. **(a)**. Tandem affinity purification was performed using HEK-293T cells stably expressing N-FLAG-S-tagged TPD52. The major hits from mass spectrometry analysis were shown in the table. **(b)**. His-tagged AMPKα1, AMPKα2, AMPKβ1, AMPKβ2 and AMPKγ1 were overexpressed and purified from *E.coli*, and GST pull-down assay using GST-TPD52 protein was carried out to determine the *in vitro* interactions. **(c)**. TPD52 interacts with AMPK endogenously in 293T cells. Irrelevant IgG was used as the immunoprecipitation control. Whole cell lysate was used as input. **(d)**. His-tagged AMPKα1 kinase domain (1-300 amino acids, AMPKαE) was overexpressed and purified from *E.coli*, and GST pull-down assay of TPD52 using the purified proteins was carried out to determine the interactions. **(e)**. Mapping the regions of TPD52 required for AMPK binding. Upper panel, schematic representation of TPD52 constructs and the minimum interaction region. Lower panel, GST-tagged TPD52 full-length (FL) or deletion mutants were purified from *E.coli*, and GST pull-down assay was performed.

### TPD52 regulates AMPK activation and cellular metabolism

Our initial observation of TPD52 interacting with AMPKα raised a question whether TPD52 can regulate AMPK activity. We found that knockdown of TPD52 by two different siRNA resulted in an increased phosphorylation of AMPKα at Ser172, but had no effect on the total AMPKα protein level in SK-BR-3 cancer cells (Figure 2(a)). On the other hand, overexpression of TPD52 in MDA-MB-231, a cell expressing low endogenous TPD52, resulted in decreased AMPK Ser172 phosphorylation, with no effect on total protein level (Figure 2(b)). These results suggest that TPD52 negatively regulates AMPKα phosphorylation, which in turn, might affect cell metabolism.

**Figure 2.**
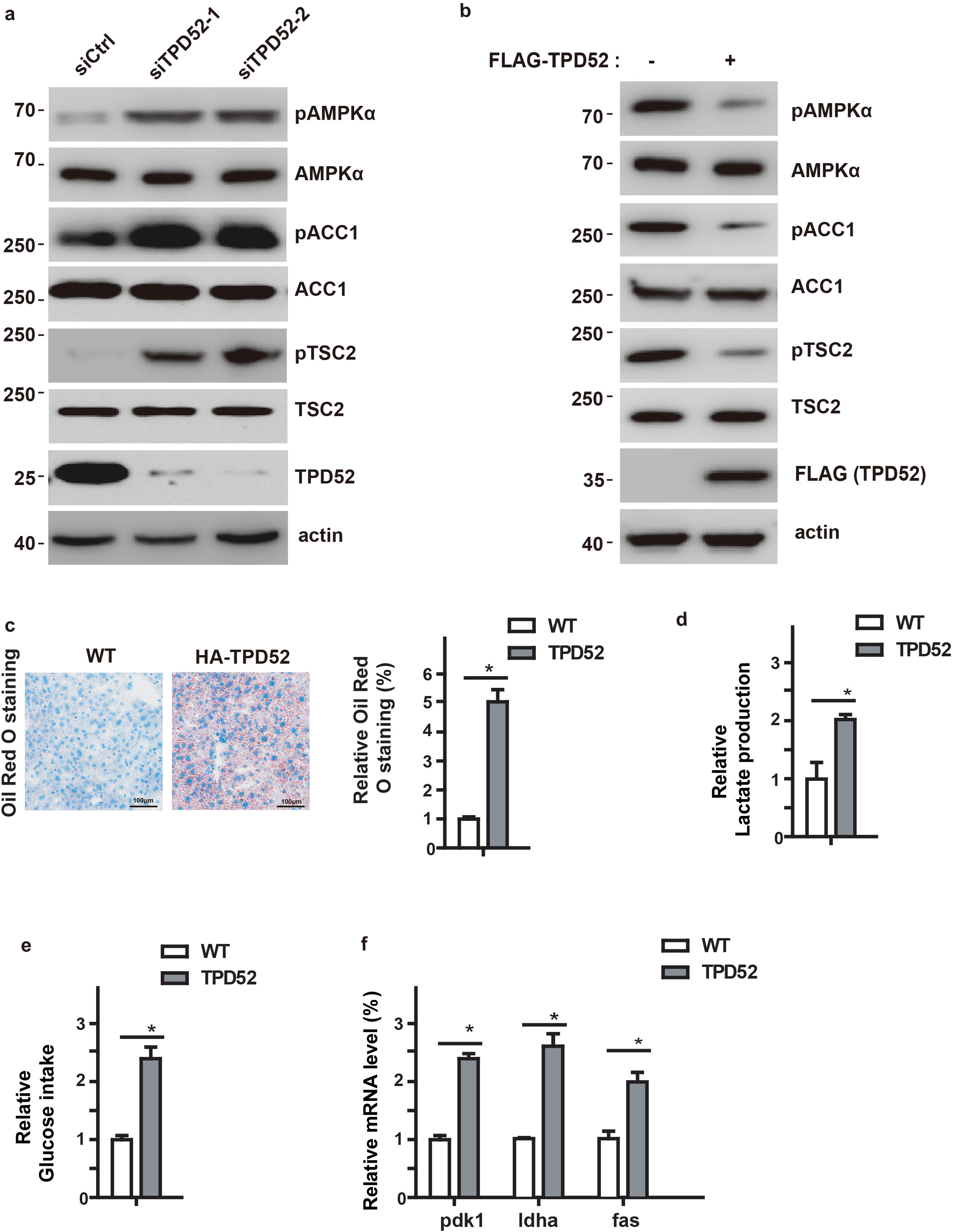
TPD52 regulates AMPK activity and cellular metabolism. **(a)**. SK-BR-3 cells were transfected with indicated siRNAs. The phosphorylations of AMPKα, ACC1 and TSC2 in cell lysates were detected by Western Blot. **(b)**. MDA-MB-231 cells were transfected with the indicated constructs, and the phosphorylations of AMPKα, ACC1 and TSC2 in cell lysates were detected by Western Blot. **(c)**. MDA-MB-231 cells were transfected with the indicated constructs, and the intracellular lipid droplet level was assessed by Oil-Red-O staining. All error bars represent the SD from the mean value of three independent experiments. *, p<0.05. **(d)**. Overexpression of TPD52 increases lactate production. The lactate production was measured in medium collected at 48 hours after transfection of control or TPD52 expressing constructs in MDA-MB-231 cells. The results represent the mean ± SEM of three independent experiments. *, p<0.05. **(e)**. Overexpression of TPD52 increases glucose intake. The glucose intake was measured in medium collected at 48 hours after transfection of control and TPD52 overexpressing constructs in MDA-MB-231 cells. The results represent the mean ± SEM of three independent experiments. *, p<0.05. **(f)**. Overexpression of TPD52 increases lipogenesis and glycolytic gene expression. Relative expression levels of *fas*, *ldha*, and *pdk1* mRNA in control or TPD52 overexpressed MDA-MB-231 cells were determined by qPCR. Transcript levels were determined relative to actin mRNA levels, and normalized to control cells. The results represent the mean ± SEM of three independent experiments. *, p<0.05.

To further confirm TPD52 regulation of AMPK activity, we examined the phosphorylation of downstream substrates of AMPK, such as ACC1 and TSC2. We found that downregulation of TPD52 significantly increased the phosphorylation of ACC1 and TSC2 (Figure 2(a)), phenomena that were consistent with increased AMPK phosphorylation. In contrast, overexpression of TPD52 decreased the phosphorylation of ACC1 and TSC2 (Figure 2(b)). These results further confirmed that TPD52 inhibits AMPK Ser172 phosphorylation and AMPKα activity.

AMPK plays important roles in cellular metabolism such as lipid metabolism and glycolysis (46–48). Previous studies showed that TPD52 expression was correlated with neutral lipid storage within cultured cells (49). However, whether this involves TPD52 regulation of AMPK by interacting with AMPK is not clear. Next, we further checked whether TPD52 regulates these metabolic processes. As shown in Figure 2(c)-2(e), TPD52 overexpression resulted in significant increases in lipid drop formation, lactate production and glucose intake. Previous studies also showed that AMPK pathway regulates expression of glycolytic genes, such as *pdk1*, *ldha* and *fas* (50). Consistently, we also found that overexpression of TPD52 increased the expression of these metabolic genes (Figure 2(f)). On the other hand, TPD52 deficiency resulted in significant decreases in lipid drop formation (Supplementary Figure S1(a)), lactate production and glucose intake (Supplementary Figure S1(b) and S1(c)). All these metabolic changes further supported that TPD52 regulates the AMPK pathway and cellular metabolism.

### TPD52 directly regulates AMPK kinase activity

As AMPK is activated by many nutrition stresses, we next examined whether these nutrition stresses might affect AMPK-TPD52 interaction. We treated cells with the established pharmacological activators of AMPK, 5-aminoimidazole-4-carboxamide-1-β-D-ribofuranoside (AICAR) or metformin, or cultured cells in amino acid or glucose deprived conditions (-AA or -Glucose). As shown in Figure 3(a), all of these treatments dramatically decreased AMPK-TPD52 interaction. These results implied that TPD52-AMPK axis is important in cell response to metabolic nutrition stresses. Next, we asked how TPD52 regulates AMPK activity. There are two possibilities for TPD52 to regulate AMPK activity. One is through the influence of the interaction between AMPK upstream kinase, LKB1 and AMPK, since AMPK activation requires Ser172 phosphorylation by LKB1. Our results in Figure 2 have already shown that TPD52 regulated AMPK Ser172 phosphorylation. To test the first possibility, we performed co-immunoprecipitation (co-IP) experiments to check whether manipulating TPD52 level might affect LKB1-AMPK interaction. As shown in Figure 3(b), overexpression of TPD52 blocked the AMPK-LKB1 interaction. Another complementary and non-exclusive possibility is that TPD52 has a direct effect on AMPK kinase activity. To test the second hypothesis, we purified ACC1 N terminal (1-200 amino acids, ACC1N) in *E.coli* as substrates for AMPK activity assay. We also purified AMPKα and TPD52, and performed the *in vitro* kinase assay. As shown in Figure 3(c), TPD52 directly inhibited AMPK phosphorylating ACC1. These results suggest that TPD52 affects AMPK activity by dual mechanisms: 1, by affecting LKB1-AMPKα interaction and in turn, AMPK Ser172 phosphorylation, 2, by directly inhibiting AMPK kinase activity toward its substrates.

**Figure 3.**
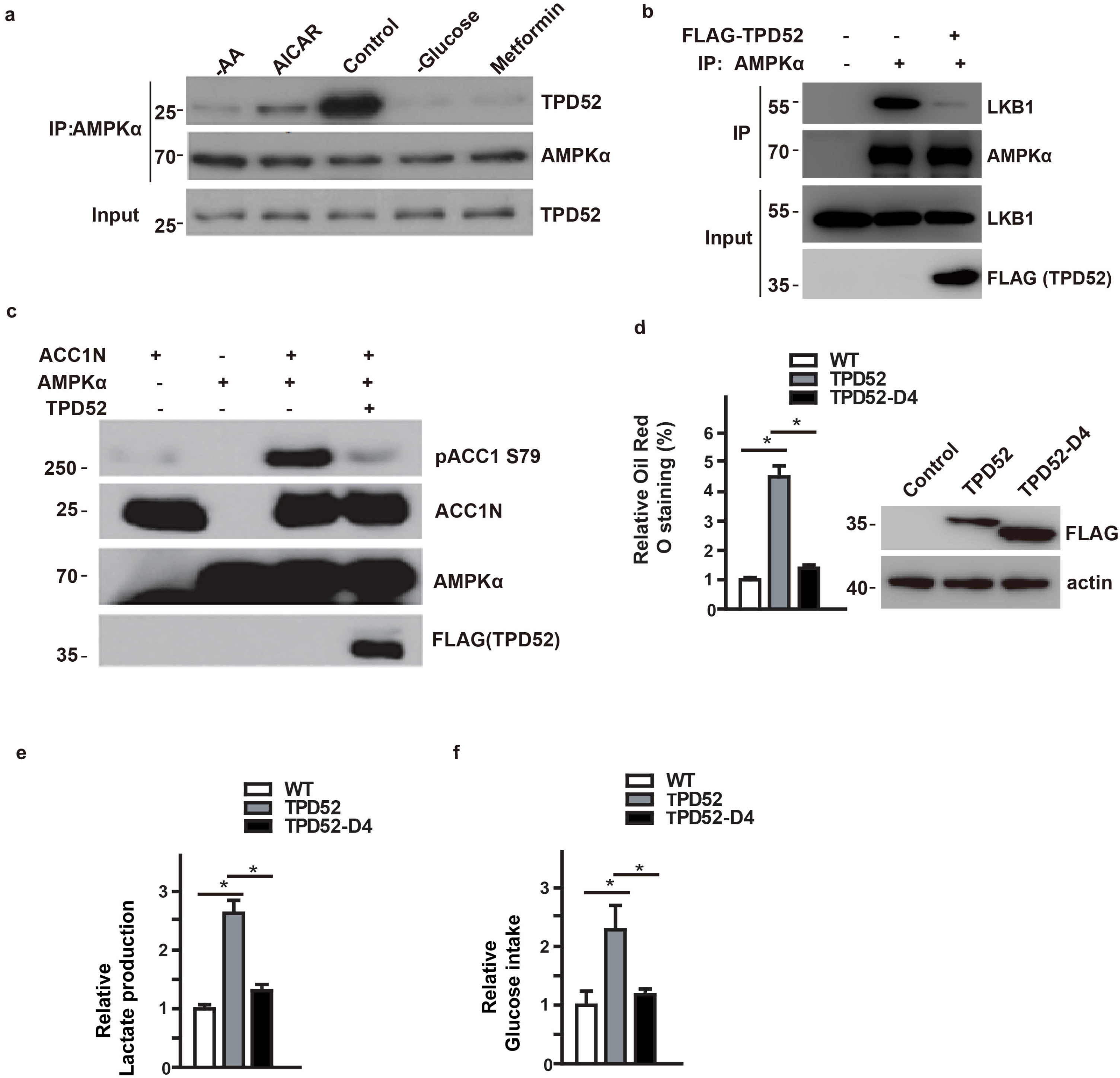
TPD52 directly regulates AMPK kinase activity. **(a)**. SK-BR-3 cells were treated as indicated for 24 hours, the interaction between TPD52 and AMPKα were examined by co-IP. **(b)**. MDA-MB-231 cells were stably transfected with TPD52, and LKB1-AMPKα interaction was examined by co-IP. **(c)**. TPD52 directly inhibits ACC1 Ser79 phosphorylation by AMPKα *in vitro*. GST-TPD52, AMPKα and ACC1 N terminal were purified from *E.coli* as indicated in Methods, and *in vitro* kinase assay was performed using ACC1 N terminal as a substrate and using AMPKα as the kinase. **(d)**. MDA-MB-231 cells were transfected with the indicated constructs, and the intracellular lipid droplets were assessed by Oil Red O staining. All error bars represent the SD from the mean value of three independent biological replicates. **(e)**. The lactate production was measured in medium collected at 48 hours after transfection of control or TPD52 WT or D4 mutant expression constructs in MDA-MB-231 cells. The results represent the mean ± SEM of three independent experiments. *, p<0.05. **(f)**. The glucose intake was measured under the same condition as previous ones. The results represent the mean ± SEM of three independent experiments. *, p<0.05.

Since TPD52 directly interacts with AMPKα through its N terminal, we hypothesized that TPD52-AMPK interaction is required for TPD52 functions in cell metabolism. As shown in Figure 3(d)-3(f), we overexpressed wild type TPD52 or N terminal deletion mutant D4 in MDA-MB-231 cells. WT TPD52, but not D4 affected lipid drop formation, lactate production and glucose intake. These results indicated that the TPD52-AMPKα interaction is required for the regulation of AMPK and its function in cell metabolism by TPD52.

### TPD52 regulates AMPK activity and cell metabolism in vivo

To explore the function of TPD52 *in vivo*, we generated *Tpd52* transgenic mice in C57BL/6J strains (Supplementary Figure S2(a)). We confirmed that the full length *Tpd52* transcript was introduced into mice by using genotyping and Western Blot (Supplementary Figure S2(b)-S2(c)). First of all, we noticed that these mice showed multiple metabolic defects, whether under nonfat diet (NFD) or high fat diet (HFD) conditions. Overexpression of TPD52 significantly increased liver lipid contents, as measured by Oil Red O staining and H&E staining, especially under the HFD condition (Figure 4(a)-4(b)). In addition, overexpression of TPD52 also resulted in increased blood glucose level under both NFD and HFD conditions (Figure 4(c)) and lower glucose tolerance (under NFD conditions, Figure 4(d)). To further clarify whether these phenotypes were related to AMPK, we measured the activation status of AMPK in the liver tissues. Consistent with our cell based assay results, overexpression of TPD52 in livers resulted in a drastic decrease of phosphorylation of AMPKα (p-AMPKα) as well as precursor SREBP1 (SREBP1-p), and an increase of nuclear processed SREBP1 (SREBP1-n) (Figure 4(e)). We also abserved an obvious decrease of phosphorylation of ACC1 and AMPKα (Figure 4(f)) in the livers from the TPD52 transgenic mice, under NFD and HFD conditions, indicating that AMPK is less activated in *Tpd52* overexpressing mice. At the gene transcription level, expression levels of glycolytic genes, such as *scd1*, *fas*, *pdk1*and *ldha* were significantly increased in TPD52 overexpressing mice (Figure 4(g)-4(h)). These phenotypes were consistent with the function of AMPK in glycolysis and lipogenesis (47,48).

**Figure 4.**
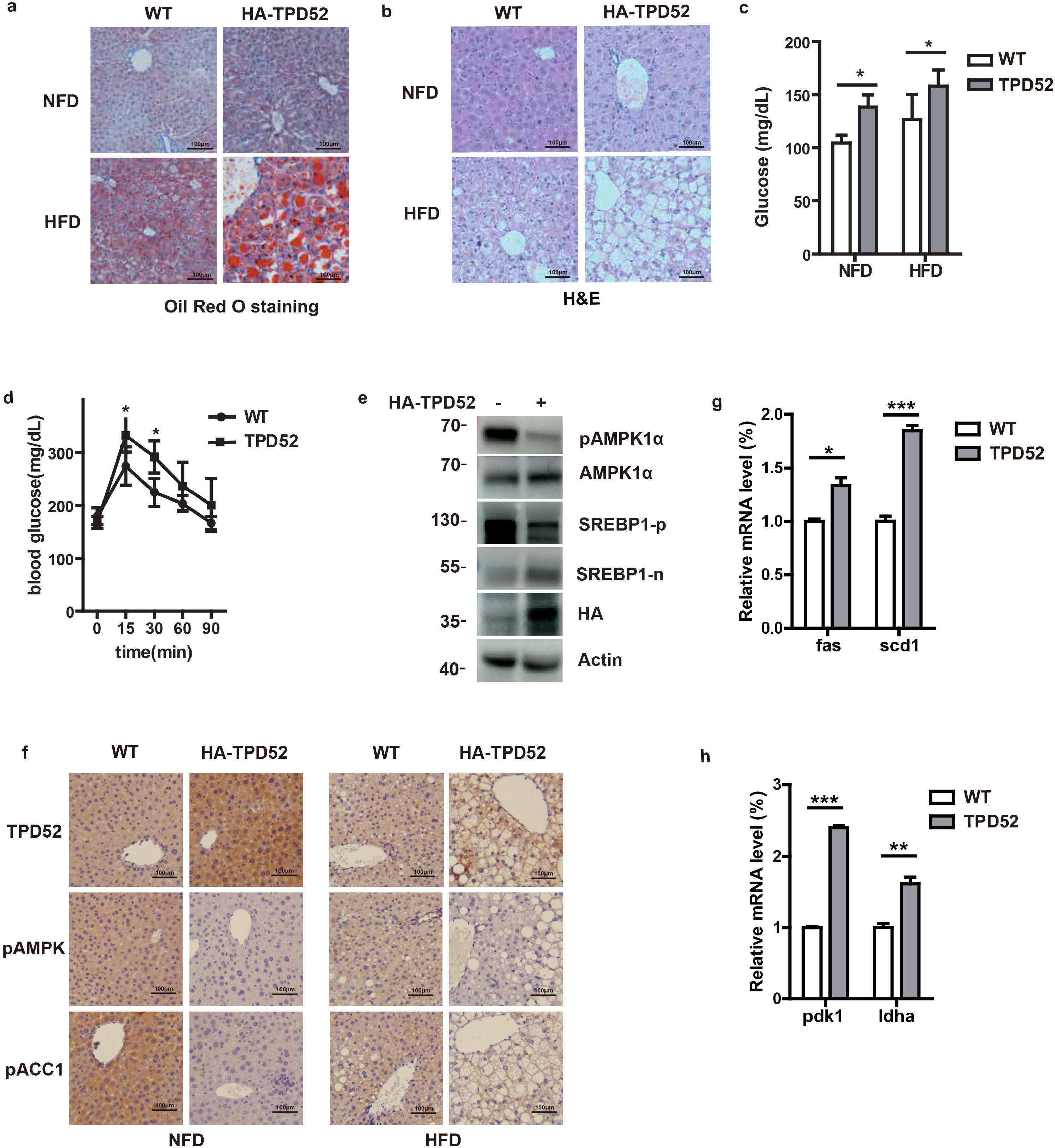
TPD52 regulates AMPK activity and cell metabolism *in vivo*. **(a). and (b)**. The Oil Red O staining and H&E staining of the liver tissues from the NFD and HFD-fed WT and HA-TPD52 transgenic mice. **(c)**. Blood glucose levels were measured in tail vein blood samples using a glucometer. Values are expressed as mean ± SD (n = 6). *p<0.05. **(d)**. Serial changes in blood glucose levels after intraperitoneal injection of glucose in the indicated mice (n=6). Values are expressed as mean ± SD. *p<0.05. **(e)**. AMPK phosphorylation, precursor (-p) and nuclear processed (-n) SREBP1c and HA-TPD52 levels in liver tissues from the indicated mice were examined by Western Blot. **(f)**. Expression of TPD52 and phosphorylated ACC1 and AMPK in livers from the NFD and HFD-fed WT and HA-TPD52 transgenic mice was determined by immunohistochemical staining. **(g). and (h)**. mRNA levels of lipogenesis gene (*fas* and *scd1*) and glycolytic gene (*pdk1* and *ldha*) in indicated liver cells from the WT and HA-TPD52 transgenic mice were determined by qRT-PCR. Relative mRNA levels were corrected to actin mRNA levels, and normalized relative to control cells. The results represent the mean ± SEM of three independent experiments. ***p<0.001., **p<0.01., *p<0.05.

The protein expression profiles found in the Human Protein Atlas has revealed that TPD52 is upregulated in the majority of the breast cancer tissues https://www.proteinatlas.org/ENSG00000076554-TPD52/pathology/tissue/breast+cancer). To determine whether TPD52 expression correlates with patient outcome, we analyzed the overall survival (OS) with a TCGA-BRCA cohort of 1109 Breast invasive carcinoma samples (https://portal.gdc.cancer.gov,TCGA-BRCA, V13.0, 2018). Indeed, the log-rank test demonstrated that the overall survival for patients with low TPD52 expression in tumor tissues was significantly higher than those with high TPD52 expression (P=0.0036, Figure 5(a)), suggesting that TPD52 expression correlates with poor patient prognosis. To explore if the level of TPD52 expression has any association with the metabolic status in tumors, we also used the RNA-seq dataset of gene expression profiles of the above TCGA-BRCA consortium to analyse the co-expression profiles between TPD52 and the metabolic genes. We observed that the expression profiles of TPD52 and the lipogenesis genes such as FAS, SCD1, ACACA/ACC1 were significantly correlated (R=0.274, 0.386, 0.358, respectively, Figure 5(b)), which is consistent with our *in vivo* data from mice. We also found that the expression profiles of TPD52 and the AMPK downstream glycolysis genes such as Aldoa and Ldha had a positive correlation (R=0.169, 0.146, respectively, Figure 5(c)), indicating that there may be a negative correlation between TPD52 expression level and AMPK activity.

**Figure 5.**
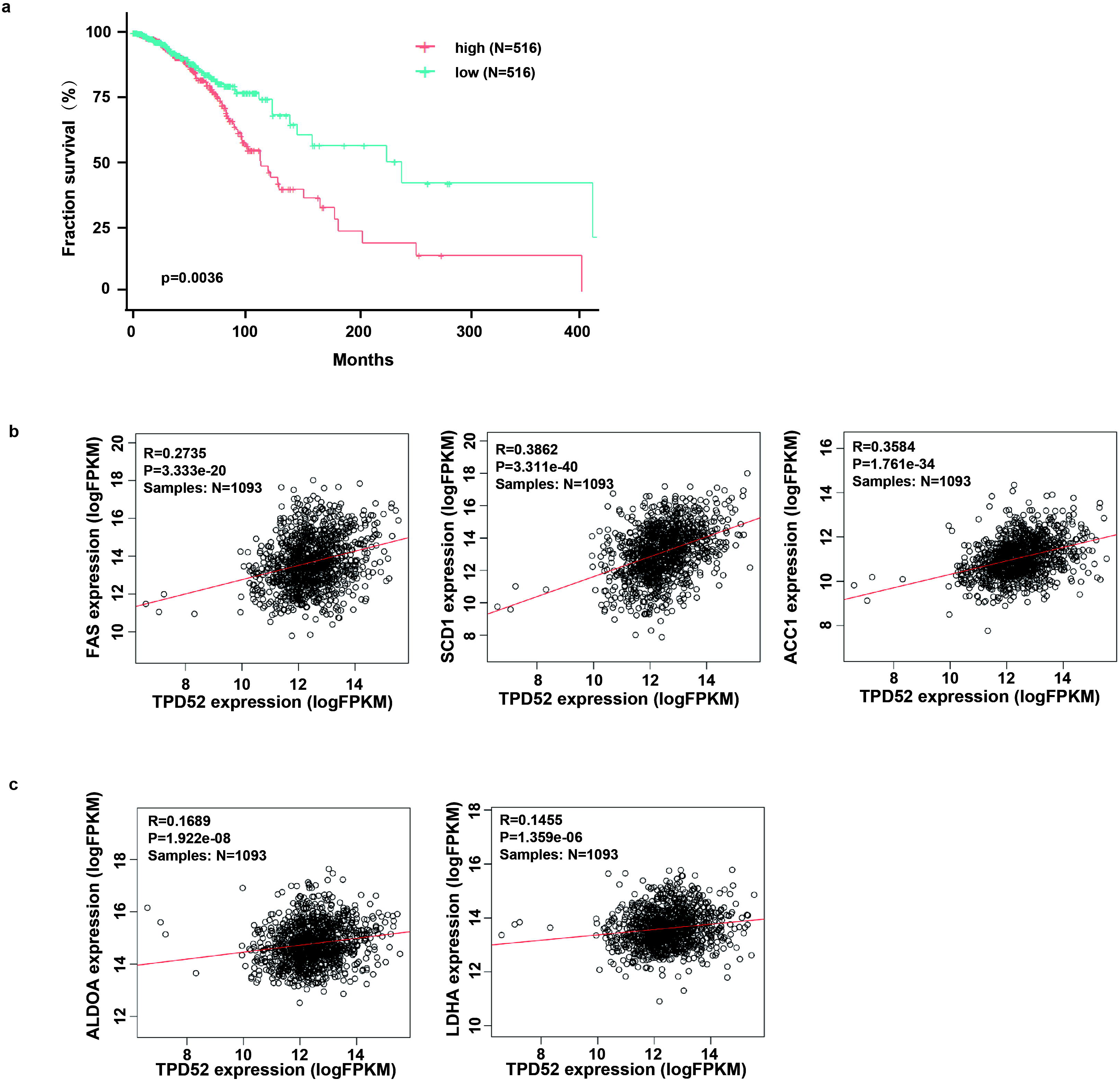
TPD52 expression is associated with clinical outcome and metabolic genes expression in breast cancer patients. **(a)**. Kaplan-Meier (KM) survival analysis of the breast invasive carcinomas patient in TCGA cohort. The patients were divided into high- and low- expression group based on the medium FPKM cutoff value of TPD52 level. **, P < 0.01. **(b)**. TPD52 and the lipogenesis genes (FAS, SCD1 and ACACA/ACC1) are co-expressed in breast invasive carcinomas. Correlation was calculated using Pearson’s correlation coefficients (R). **(c)**. TPD52 and the glycolysis genes (Aldoa, Ldha) are co-expressed in breast invasive carcinomas. Correlation was calculated using Pearson’s correlation coefficients (R).

Taken together, our results suggest that TPD52 plays a key role in AMPK activation and controls the lipid and glucose metabolism *in vivo*.

## Discussion

The LKB1-AMPK pathway has been identified as a central regulator of energy metabolism, and dysregulation of this pathway has been implicated in many cancers, including breast cancer (31–33). However, other than LKB1 mutations, how this pathway might be dysregulated in breast cancer remains unclear. In this study, we have established TPD52 as a regulator of energy stress-induced AMPK activation and cell metabolism. TPD52 transgenic mice also showed multiple metabolic defects, which mimics many phenotypes observed with AMPK knockout mice (51). AMPK can also be activated in response to calcium flux through CaMKKβ (15–17). Therefore, it would be interesting to test whether TPD52 also functions in a similar way in response to calcium flux in future studies.

TPD52 directly bound to AMPKα, but not LKB1 (Fig. 1 and data not shown). Nutrition or energy stress disrupted AMPK-TPD52 interaction. It is possible that TPD52 and LKB1 might bind to the same regions of the kinase catalytic domain of AMPKα coordinately to regulate AMPK phosphorylation. On the other hand, TPD52 also directly regulates AMPK phosphorylating ACC1 *in vitro*. The kinase domain of AMPKα conformation change is very important for AMPK activation (9,10). TPD52 binding might directly influence AMPKα conformation and in turn, affect its kinase activity.

*Tpd52* (tumor protein D52), belongs to the human *Tpd52* gene family that is made up of four genes *hD52*, *hD53*, *hD54* and *hD55*. The *Tpd52* gene (*hD52*) maps to chromosome 8q21 and is known to be involved in the regulation of vesicle trafficking and exocytotic secretion (36–38). In addition, the 8q21 locus containing the *Tpd52* gene is frequently amplified in tumors, including breast and liver. Thus far, only one study showed that overexpression of murine TPD52 can initiate cellular transformation, tumorigenesis and progression to metastasis (52). Here we found that TPD52 regulates AMPK pathway and cell metabolism in cells and in mice. By exploring TCGA-BRCA cohort data, we also found that TPD52 expression is associated with clinical outcome and metabolic gene expression in breast cancer patients. As cell metabolism dysregulation is a key character for tumor initiation and progressing. Our studies provided new insights for TPD52 associated cancer etiology.

Our findings will have a significant impact on the dissection of components in the pathway controlling AMPK activity. Furthermore, since AMPK plays important roles in a number of cell signaling pathways such as cell growth, autophagy in addition to cell metabolism (24,25). The key role of AMPK places it as an ideal therapeutic target for the treatment of obesity, insulin resistance, type 2 diabetes and cancer (53,54). Our findings may also have important implications for multiple disease etiology and response to therapy.

## Acknowledgements

We thank the Laboratory Animal Platform of National Center for Protein Sciences-Beijing (NCPSB) for assistance with animal studies. We thank Dr. Yang Dong and Gao Chao for their kind help in bioinformatics analysis.

