## supplementary file for "Tumor protein D52 (TPD52) affects cancer cell metabolism by negatively regulating AMPK"

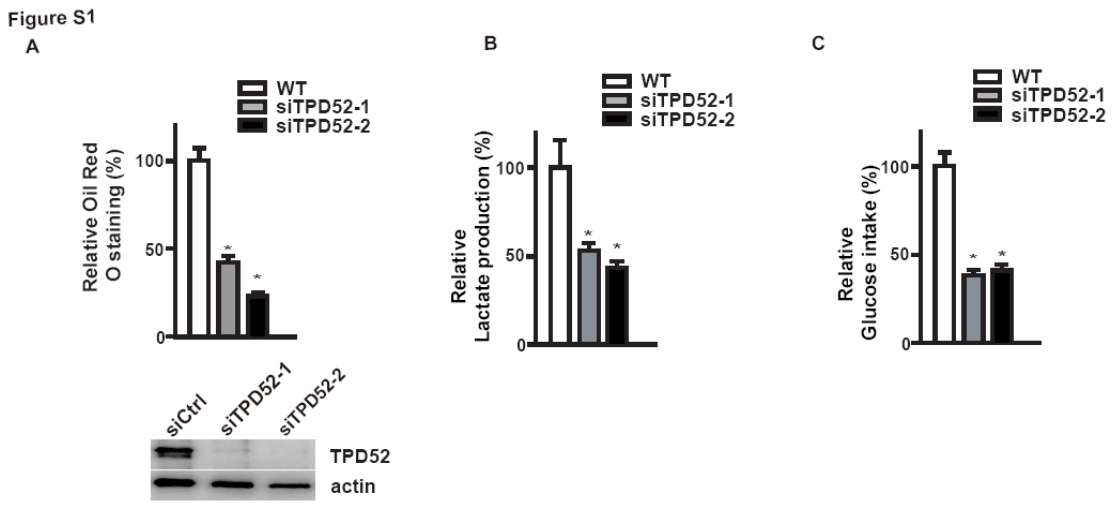

**Supplementary Figure 1. TPD52 regulates cellular metabolism in T-47D cells.**

**A-C.** T-47D cells were transfected with the indicated siRNAs, and the intracellular lipid droplet (A), the lactate production (B) and the glucose intake (C) were measured in medium collected at 48 hr after incubation of control and TPD52 knocked down T-47D cells. The results represent the mean ± SEM of three independent experiments. *, p<0.05.

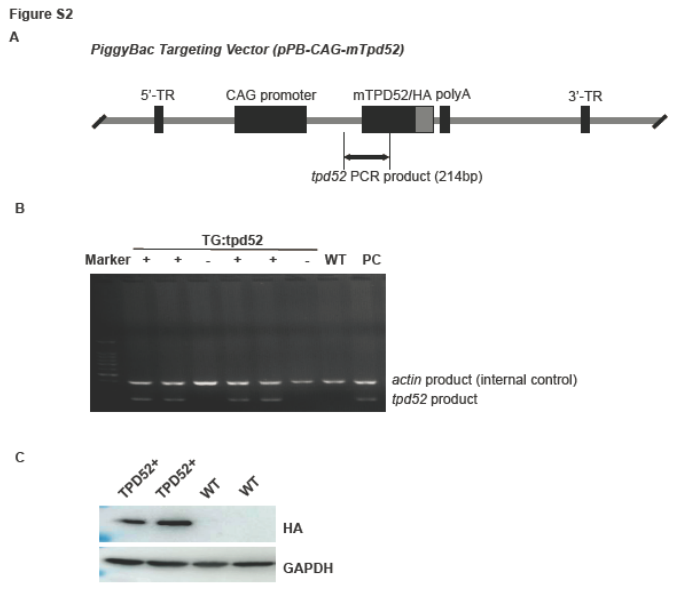

**Supplementary Figure 2. Generating *Tpd52* transgenic mice.**

**A.** Schematic figure of the PiggyBac transposon gene expression vector for mouse *Tpd52* gene.

**B.** Representive of genotyping results of the transgenic (TG: *tpd52*) mice.

**C.** Western Blot detection of exogenous HA-mTPD52 expression in the transgenic mice.
